# Decoding DNA methylation and non-coding RNAs mediated regulatory landscape of breast tumor and adjacent normal tissues

**DOI:** 10.1101/2025.02.17.638770

**Authors:** Swapnil Kumar, Vaibhav Vindal

**Affiliations:** Department of Biotechnology & Bioinformatics, School of Life Sciences, University of Hyderabad, Hyderabad, 500046, India.

**Keywords:** Tumor-adjacent normal tissues, breast cancer subtypes, multilayer network, integrative network, epigenetic regulation, long non-coding RNA, microRNA

## Abstract

Tumorigenesis not only involves perturbations of mRNAs but also microRNAs (miRNAs), long non-coding RNAs (lncRNAs), and DNA methylation (DNAm). Additionally, the tumor-adjacent normal tissues (TANTs) that are used as controls in cancer studies are not completely normal and, if explored systematically, can help to understand tumorigenesis better. In this regard, expression and DNAm profiles of breast tumor tissues (viz., Luminal-A, Luminal-B, Her2+, Basal-like, and Normal-like) and TANTs were used to construct and analyze their integrative networks. These networks included regulations among genes, transcription factors, lncRNAs, and miRNAs, along with gene/lncRNA methylations. This followed the identification of candidate gene *PCLAF* having transitivity value one across all six tissue networks. Further, 503 common hubs (388 protein-coding, 100 miRNAs, and 15 lncRNAs), including seven methylated genes (*SPI1*, *STAT5A*, *RORC*, *SPDEF*, *ELF5*, *HOXB2*, and *RUNX3*), and 149 common bottlenecks (145 protein-coding, three miRNAs, and one lncRNA), including three methylated genes (*SPI1*, *SPDEF*, and *RUNX3*) and one methylated lncRNA (*PART1*), were identified. Moreover, 145 nodes, including three methylated genes (*SPI1*, *SPDEF*, and *RUNX3*) and three miRNAs (hsa-miR-34a, hsa-miR-149, and hsa-miR-141), were identified as both hubs and bottlenecks (HBs) across all six tissues. However, none of the lncRNAs were found HBs therein. Genes and lncRNAs with higher degrees, betweenness, eigenvector centrality, and pagerank were found more preferably methylated than others. Further, pathway analysis of hubs and bottlenecks across all six tissues revealed their significant involvement in the pathway of Transcriptional misregulation in cancer. Overall, these findings uncover new insights into the epigenetic mechanisms of tumorigenesis.

## Introduction

Breast cancer is the major cause of death and also the most frequently detected cancer in women worldwide [Bray et al., 2024]. It is of a heterogeneous nature with multiple subtypes and highly complex underlying mechanisms that develop in the mammary tissues, including lobules and ducts. Many of the complex signaling pathways involved in normal human breast development, predominantly HER2 signaling, estrogen receptor (ER) signaling, and canonical Wnt signaling, are dysregulated or hijacked in case of the diseased condition. The epigenetic and genetic alterations within cancer cells enable them to escape the normal mechanisms governing their death, survival, division, proliferation, and migration [Sever and Brugge 2015]. Further, any alterations that lead to the activation of proto-oncogenes can result in dysregulation and hyperactivation of associated signaling pathways. Similarly, any alterations inactivating tumor suppressors can lead to the elimination of critical negative regulators of associated signaling pathways that are responsible for their proper functionalities.

Cancer studies, including clinical and other experimental studies, generally utilize tissue samples from human patients or mouse models of cancer. For example, breast cancer studies mainly involve experimental exploration of breast tumor tissues (BTTs) compared with tumor-adjacent normal tissues (TANTs) as controls. Notwithstanding, these TANTs are not completely normal and harbor mutations and other abnormalities linked to cancer. Thus, the TANTs provide an intermediate state between healthy and tumor tissues [Slaughter et al. 1953; Heaphy et al. 2009; Aran et al. 2017]. The exploration of this intermediate state might provide key insights into the pre-cancerous mechanisms that lead to the transformation of tissues from a healthy normal state to a tumor state. Despite the known tumorigenic features of TANTs, epigenetic regulations in these tissues have not been explored yet. The epigenetic regulation is a heritable mechanism that involves gene expression regulation without altering their actual nucleotide sequences and includes regulations caused by DNA methylation (DNAm), chromatin remodeling, histone modification, and RNA-mediated gene targeting [Holliday, 1994]. The expression of genes is regulated epigenetically by microRNAs (miRNAs), long non-coding RNAs (lncRNAs), and DNAm, and any alteration of epigenetic regulations in molecular and cellular pathways can lead to a diseased condition [Dawson and Kouzarides, 2012]. The role of epigenetic alterations, i.e., regulations of gene expressions by molecular entities such as lncRNAs, miRNAs, and DNAm at the pre- or post-transcriptional levels, has become a fascinating area. Further, developing a single disease model by incorporating all this information is computationally intensive and challenging; however, if modeled, it can aid in the comprehensive understanding of underlying mechanisms.

LncRNAs are the longest endogenous non-coding RNAs (ncRNAs) with approximate lengths from 200 nucleotides (nt) to 100 kb. These ncRNAs are key epigenetic modifiers and play a crucial role in breast cancer progression and metastasis by degrading or silencing target mRNAs, sponging miRNAs, or else targeting the machinery involved in the miRNA biogenesis. For example, *ANCR*, a tumor suppressor lncRNA, can inhibit cell migration and metastasis in breast cancer by decreasing the expression of *RUNX2* [Li et al., 2017]. Further, miRNAs are small ncRNA molecules with approximate lengths of 18-21 nt and regulate the expressions of target genes post-transcriptionally either by silencing or degrading them. It has been reported that miR-373-3p and miR-206 regulate the expression of *ITGA2* in breast cancer [Ding et al., 2015; Adorno-Cruz et al., 2021]. Moreover, DNAm involves attaching methyl group (-CH_3_) to cytosine with the help of three types of DNA methyltransferases (DNMTs), viz., DNMT1, DNMT3A, and DNMT3B, without altering the DNA sequence [Okano et al., 1999; Feng and Lou, 2019]. Any changes to the normal methylation pattern of genes lead to abnormal methylation, which includes either hypermethylation or hypomethylation [Kanwal and Gupta, 2012]. Several studies have reported the association of abnormal methylation with the causes of cancers, including breast cancer [Prajzendanc et al., 2020; Nunes et al., 2020; Hong & Kim, 2021; Bévant et al., 2021; Chiappinelli & Baylin, 2022]. It has been reported that the overexpressed miR-148a/miR-152 in breast cancer suppressed cell proliferation, migration, colony formation, and angiogenesis; however, the aberrantly upregulated *DNMT1* was inversely correlated with miR-152/miR-148a expression [Xu et al., 2013]. Moreover, one other study reported that the methylation at the promoter of *MEG3* played a key role through the miR-494-3p/OTUD4 axis in breast cancer growth [Zhu et al., 2022]. LncRNAs, miRNAs, and DNAm epigenetically coordinate to provide precise expressions of genes and thus present exciting therapeutic options.

In the present study, we have focused on the ncRNAs and DNAm-related epigenetic regulations in TANTs and breast cancer subtypes. For this, six (five subtypes and one TANT) epigenetic-mediated multilayer regulatory networks were constructed by utilizing gene expression and DNAm profile data coupled with prior information of transcription factors (TFs)-targets, lncRNAs-targets, and miRNAs-targets as retrieved from various sources. Subsequently, these networks were analyzed by exploring various topological properties, such as degree, betweenness, and clustering coefficient of each regulatory entity participating therein. Besides, regulatory patterns of methylated and non-methylated genes and lncRNAs were also studied. This is the first extensive study exploring epigenetic-mediated multilayer networks of TANTs along with different subtypes of breast cancer, which resulted in the revelation of some novel and key molecular players (protein-coding genes, miRNAs, lncRNAs, and methylation patterns) across TANTs and BTTs responsible for breast cancer tumorigenesis. It also revealed the distinctive regulatory patterns of methylated genes and lncRNAs compared to the non-methylated ones. However, these identified molecules need to be investigated further using *in vitro* and *in vivo* experiments to better understand the epigenetics of breast tumors and also to develop more effective therapeutic and diagnostic methods for the early stage of the disease.

## Methods

### Acquisition and processing of breast cancer data

Four different types of data, i.e., the gene expression, miRNA expression, lncRNA expression, and DNA methylation data of BTTs and TANTs, were downloaded from the database of TCGA (The Cancer Genome Atlas) [Tomczak et al. 2015; Weinstein et al. 2013] using the TCGAbiolinks Bioconductor/R package [Colaprico et al. 2016]. Only 1057 BTT samples and 93 TANT samples with all four data types were considered for further analysis.

These four data sets of BTTs were further stratified into five data sets for five different breast cancer subtypes, viz., Basal, Her2, LumA, LumB, and NormL. Thus, there were four data types, viz., gene expression, lncRNA expression, miRNA expression, and DNA methylation for each of the six tissue types, including BTT and TANT. For further analysis, both the gene and lncRNA expression profiles were normalized using the TCGAbiolinks package. Subsequently, from all three expression data (viz., gene, lncRNA, and miRNA), only the expression data of protein-coding genes, lncRNAs, and miRNAs having an average value of one count per million (CPM) across all the samples were considered for the current study. The CPM values for each protein-coding gene, lncRNA, and miRNA were calculated using the “cpm” function of the edgeR package [Robinson et al. 2010]. Furthermore, the DNA methylation (DNAm) data contained some cross-reactive probes and polymorphism-mapped probes. Besides this, it also contained missing values. Thus, the DNAm data were, first, processed by imputing all the missing values using the “impute.knn” function with default parameters from the impute R package, which implements the k-nearest neighbor method for the missing value imputation. Subsequently, all the cross-reactive probes and probes mapping to polymorphism were filtered out before further analysis.

### Target prediction of miRNAs and lncRNAs

All miRNAs and lncRNAs with one CPM in each tissue type were subjected to the identification of their targets using different databases. Targets of miRNAs were identified using RegNetwork [Liu et al. 2015], miRanda [John et al. 2004], miRTarBase [Hsu et al. 2011], and TargetScan [Agarwal et al. 2015]. Meanwhile, targets of lncRNAs were retrieved from LncTarD v2.0 [Zhao et al. 2023] and LncRNA2Target v3.0 [Cheng et al. 2019]. Further, for each tissue type, the Pearson correlation coefficient values between all possible pairs of miRNA-gene, miRNA-lncRNA, lncRNA-gene, and lncRNA-miRNA were calculated using the expression data of protein-coding genes, lncRNAs, and miRNAs with minimum one CPM expression value. Subsequently, all pairs of miRNA-gene, miRNA-lncRNA, lncRNA-gene, and lncRNA-miRNA with negative correlations were mapped to the previously identified pairs of miRNA-target and lncRNA-target. This resulted in the identification of condition-specific pairs of miRNA-gene, miRNA-lncRNA, lncRNA-gene, and lncRNA-miRNA, which further will be used to construct the integrative networks of different tissues, including subtypes and TANTs.

### Target prediction of transcription factors

Pairs of TFs and their target genes for each tissue type were predicted with the help of pandaR package [Schlauch et al. 2017] using three prior data, including gene expression profiles of corresponding tissues, protein-protein interactions, and TFs-motif information. The PPI data were retrieved from the STRING [Szklarczyk et al. 2023] and CancerNet [Meng et al. 2015] databases, whereas the TF-motif information was from the web page of pandaR. After prediction, TF-target pairs were filtered for further analysis based on three criteria: (a) should have a score greater than zero, (b) should have a motif, and (c) TFs and target genes both should have a CPM value of one.

### Prediction of significantly methylated genes

Most of the genes are methylated mainly at their promoter region; however, some genes have also been reported to be methylated at their body region or promoter as well as body region. Further, methylation and gene expression are mainly negatively correlated; however, they can be positively correlated too. In the present study, for each tissue type, all genes, including protein-coding and lncRNAs with significant methylation either at the body or promoter regions, were predicted using a customized R script. In this, all known probe-gene pairs as per the Illumina Infinium HM450K manifest file were subjected to the calculation of Pearson correlation coefficients along with corresponding P-values and adjusted P-values between the β-value of methylation probe and the expression value of the respective gene. This led to the detection of significant probe-gene pairs. The P-value was adjusted using the BH (Benjamini & Hochberg) method, and the significance level was considered at an adjusted P-value of less than 0.05.

### Construction of integrative networks

All the previously identified miRNA-gene, miRNA-lncRNA, lncRNA-gene, lncRNA-miRNA, and TF-gene pairs, along with methylation information of genes and lncRNAs, were integrated to construct a whole-genome integrative network for each subtype of breast cancer and ANTs. Thus, a whole-genome integrative network for each tissue type was constructed. However, these networks were highly complex for further analysis. This can be due to too many regulatory connections among molecules, such as TFs, genes, and miRNAs.

### Estimation of genes, miRNAs, and lncRNAs with high centralities

The complete network of each tissue type (viz. Basal, Her2, LumA, LumB, NormL, and TANT) was analyzed to identify genes, miRNAs, and lncRNAs with high degree, high betweenness, and clustering coefficient 1 (CC = 1). The nodes with high degree and betweenness values were considered hubs and bottlenecks and were identified using the Pareto-principle of 80:20, where the average degree and betweenness values of the top 20% of nodes were taken as a cutoff to decide the nodes with high degree and betweenness value. Further, these networks may contain some false positive regulations, especially between TFs and genes, and hence may affect the values of degree and betweenness, leading to the biased identification of hubs and bottlenecks. To reduce the chance of any biases, each network was reconstructed by rewiring 5% of its total edges, which followed the calculation of the degree and betweenness of the resultant network and then the identification of hubs and bottlenecks. This procedure was repeated 20 times for all six networks, each time by rewiring a new set of 5% edges that had not been rewired previously. Subsequently, only nodes that were hubs or bottlenecks in the original network of each tissue and also across all the corresponding 20 rewired networks were considered true hubs and bottlenecks for further analysis.

Degree: It is defined for a node as the number of all nodes connected directly with that node. Highly connected nodes in networks are called hubs and play vital roles in their functional and behavioral organizations (Albert et al. 2000; Carlson et al. 2006; Han et al. 2004; Jeong et al. 2001).

Betweenness centrality: It is defined for a node as the number of shortest paths among all pairs of nodes crossing that node divided by the number of shortest paths among all pairs of nodes present in the network (Freeman 1977). Nodes with high betweenness values are called bottlenecks and are considered vital for the information flow from one part to another part of the network.

Clustering coefficient: It is defined for a node as the number of triangles that the node has with its neighbors divided by the number of all feasible triangles that the node can form (Ravasz et al., 2002; Watts & Strogatz, 1998). It is also known as Transitivity and provides clues about the inter-connectedness of every node with its neighboring nodes in a given network and thus reveals the modularity of the subnetworks present therein.

### Topologies of methylated and non-methylated genes

We also explored methylated and non-methylated genes with the help of their regulatory patterns in the integrative networks and their involvement in different pathways. For this, all genes, including protein-coding and lncRNAs, having significant correlations between their expression and β-values, as inferred in the previous step, were considered methylated genes. The rest of the genes, including genes without significant correlations between their expression and β-values, were considered non-methylated. Further, we investigated seven different topological properties of these genes, viz., degree, betweenness, transitivity, closeness centrality, eigenvector centrality, and pagerank.

Closeness centrality: It is defined for a node as the reciprocal value of the sum of all distances between that node and all other nodes in a given network (Freeman, 1978). Thus, it measures the number of required steps to access all the other nodes from a node of interest. In other words, it indicates the extent of closeness of every node in a network with other nodes present therein.

Eigenvector centrality: It is defined as the eigenvector value of the largest eigenvalue of a given network’s adjacency matrix (Bonacich 1972; Ruhnau 2000). It provides the measurement of the influence a node has in a given network. If a node of interest is connected to nodes having high scores, these high-scoring nodes have higher contributions to that node’s score compared to the low-scoring nodes. Thus, genes with high eigenvector scores reveal that these genes are connected with other genes that have high scores, and hence, it refers to the relative importance of every node in a given network due to their interactions with other nodes present therein.

PageRank: If node A is directly connected with nodes V_1_, V_2_, …, and V_n_ by edges pointing to them. The PageRank of node A can be defined as PR(A):

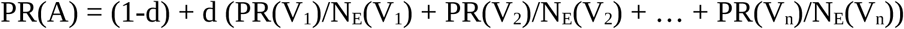

Where d is a damping factor with a value ranging between 0 and 1, and as default, it is set to 0.85. Further, N_E_(V_1_), N_E_(V_2_), …, and N_E_(V_n_) are the numbers of edges going out of nodes V_1_, V_2_, …, and V_n_, respectively [Brin & Page, 1998].

### Over-represented pathway analysis

All the hub and bottleneck genes across all six tissue types were analyzed for their involvement in various kinds of KEGG (Kyoto Encyclopedia of Genes and Genomes) pathways, viz., Cellular Processes (viz. Cell growth and death, Cellular community, Cell motility, etc.), Environmental Information Processing (viz., Signaling molecules and interaction, Membrane transport, and Signal transduction), Genetic Information Processing (viz., Transcription, Translation, Replication and repair, etc.), Human Diseases (viz., Cancers, Cardiovascular disease, Immune disease, Infectious disease, etc.), and Organismal Systems (viz., Endocrine system, Immune system, Nervous system, etc.) using the clusterProfiler package. For this, each KEGG term with adj P-value < 0.05 was only considered significantly over-represented across all six tissue types.

## Results

### Breast tumor and adjacent normal tissues

The gene expression data contained 56602 genes, including 19054 protein-coding genes and 5417 lncRNAs as per the HGNC (Human Gene Nomenclature Committee) data. Further, the miRNA expression data contained 1881 miRNAs, and the DNA methylation data included methylation levels of 25978 genes, including protein-coding genes and lncRNAs. Of 1057 samples, 183 were of Basal, 81 were of Her2, 553 were of LumA, 200 were of LumB, and 40 were of NormL. Besides, there were 93 samples of TANT.

Gene and lncRNA expression data were normalized for further analysis, as discussed in the Methods section, which followed the filtration of lowly expressed genes/lncRNAs. Any gene or lncRNA with a CPM value less than one was considered lowly expressed and non-informative and hence filtered out from the data. Similarly, the miRNA expression data were also preprocessed by filtering out all miRNAs with less than one CPM. The distribution of samples along with protein-coding genes, miRNAs, and lncRNAs with CPM values higher than one across all six tissue types (five subtypes and one TANT) are displayed in Table 1.

**Table 1.** Distribution of samples along with genes (protein-coding), miRNAs, and lncRNAs with CPM values higher than one across all six tissue types.

| Tissue | Sample | Gene | miRNA | lncRNA |
| --- | --- | --- | --- | --- |
| Basal | 183 | 13755 | 460 | 3834 |
| Her2 | 81 | 13574 | 413 | 3745 |
| LumA | 553 | 13522 | 385 | 3579 |
| LumB | 200 | 13536 | 417 | 3663 |
| NormL | 40 | 13636 | 390 | 3599 |
| TANT | 93 | 13547 | 380 | 3497 |

### TFs, miRNAs, lncRNAs, and their targets

The targets of TFs from the previously filtered gene list across all tissue types were predicted based on the expression profiles of genes with prior information on PPIs and TFs-motifs, as discussed in the Methods section. Similarly, the targets of miRNAs and lncRNAs from the previously filtered respective lists across all tissue types were predicted based on the expression profiles of genes, miRNAs, and lncRNAs with prior information on miRNA-target and lncRNA-target, as discussed in the Methods section. The detailed distribution of TF-target, miRNA-target, and lncRNA-target across all five subtypes and TANTs is available in Table 2.

**Table 2.** Distribution of targets of TFs, miRNAs, and lncRNAs across all six tissue types.

| Tissue | TF-Gene | miRNA-Gene | miRNA-lncRNA | lncRNA-miRNA | lncRNA-Gene |
| --- | --- | --- | --- | --- | --- |
| Basal | 3114127 | 274370 | 180 | 293 | 1708 |
| Her2 | 3077918 | 260350 | 181 | 280 | 1782 |
| LumA | 3066011 | 253884 | 159 | 286 | 1503 |
| LumB | 3068670 | 255988 | 177 | 302 | 1553 |
| NormL | 3094582 | 257214 | 172 | 303 | 1738 |
| TANT | 3073914 | 242995 | 168 | 277 | 1629 |

TF, Transcription factor.

### Methylated genes and lncRNAs

Significantly methylated genes and lncRNAs were identified across all tissue types, as discussed in the Methods section. If a probe-gene or probe-lncRNA pair had a positive or negative correlation (between the β-value of the probe and the expression value of the gene/lncRNA) with an adjusted P-value less than 0.05, it was considered a significantly methylated gene/lncRNA. The distribution of all significantly methylated regions, including body and promoter regions, of genes and lncRNAs across all six tissue types is available in Table 3 and Fig 1.

**Table 3.** Distribution of significantly methylated regions of genes and lncRNAs across different tissue types.

| Tissue | MBR-Gene | MPR-Gene | MBR-lncRNA | MPR-lncRNA |
| --- | --- | --- | --- | --- |
| Basal | 1912 | 3224 | 18 | 22 |
| Her2 | 1005 | 1557 | 14 | 16 |
| LumA | 2973 | 5108 | 32 | 15 |
| LumB | 2238 | 3663 | 19 | 16 |
| NormL | 353 | 560 | 5 | 2 |
| TANT | 1390 | 2590 | 14 | 8 |

MBR-Gene, Methylated body region of a gene; MPR-Gene, Methylated promoter region of a gene; MBR-lncRNA, Methylated body region of an lncRNA; MPR-lncRNA, Methylated promoter region of an lncRNA.

**Fig 1.**
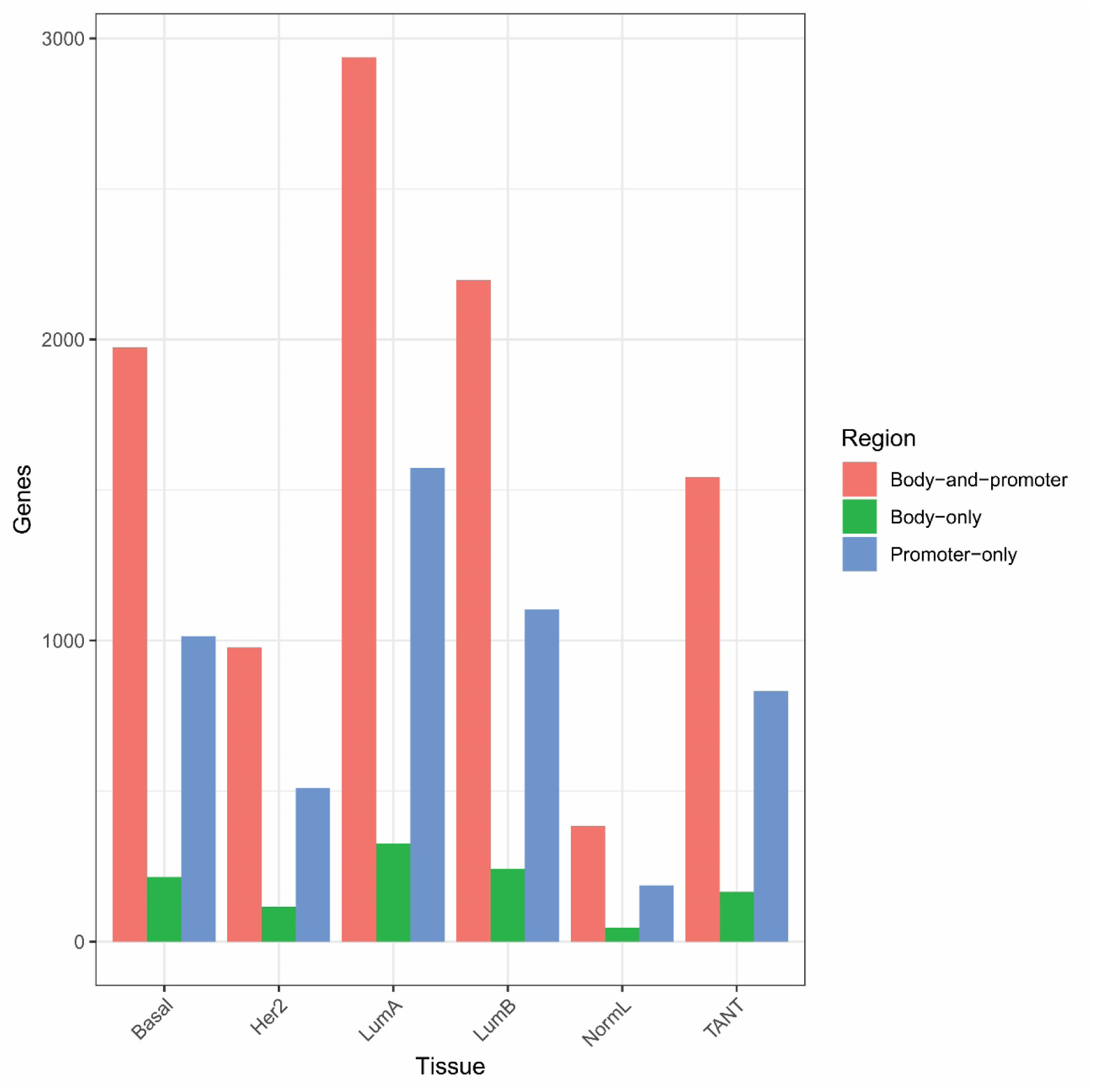
Distribution of methylated genic regions across all six tissue types

### Integrative networks of subtypes and TANTs

The integrative networks of subtypes and TANTs were constructed by combining various regulatory interactions and methylation status, as mentioned in the Methods section. These networks had two types of edges: one was the representative of regulatory interactions, connecting TFs, miRNAs, and lncRNAs to their targets, and the other was the representative of the methylation status of nodes depending on their hyper or hypomethylation. The nodes present in these integrative networks were of five types, viz., TFs, Genes, miRNAs, lncRNAs, and methylation probes. The distribution of the total number of nodes and edges in each tissue network is available in Table 4.

**Table 4.** Distribution of nodes and edges across all six tissue networks.

| Network | Nodes | Edges |
| --- | --- | --- |
| Basal | 18802 | 2557151 |
| Her2 | 16169 | 2445696 |
| LumA | 21261 | 2403969 |
| LumB | 19262 | 2413817 |
| NormL | 14631 | 2432154 |
| TANT | 17452 | 2383269 |

### Key players across subtypes and TANTs

The integrative networks of all subtypes and ANTs were analyzed to identify key molecules based on each node’s degree, betweenness, and transitivity. A node was identified as either hub or bottleneck if that node’s degree or betweenness value was above a cut-off decided by the Pareto principle of 80-20. Thus, there were 617, 580, 584, 581, 558, and 573 hubs in Basal, Her2, LumA, LumB, NormL, and TANT, respectively. Similarly, there were 354, 357, 344, 330, 347, and 322 bottlenecks in Basal, Her2, LumA, LumB, NormL, and TANT, respectively. However, these hubs and bottlenecks only included protein-coding genes and miRNAs, not any lncRNAs. Therefore, to identify lncRNAs specifically as hubs and bottlenecks, only lncRNAs participating in each network were considered, and following the same 80-20 principle, lncRNAs as hubs and bottlenecks were identified. Tables S1, S2, and S3 provide the detailed distribution of hub and bottlenecks. Further, if a gene had a CC value of one, it was also identified as a key molecule because a node with a CC value of one forms a complete sub-graph, called a clique. These cliques are the building blocks of a network. The detailed distribution of the clique-forming genes and lncRNAs (CC = 1) is available in Table 5. All the nodes with the property of being both hubs and bottlenecks and participating in clique formation are supposed to play vital roles in the organization and information flow within networks. Thus, these nodes could be considered as key players in the disease system that play vital roles in the disease’s development and progression.

**Table 5.**
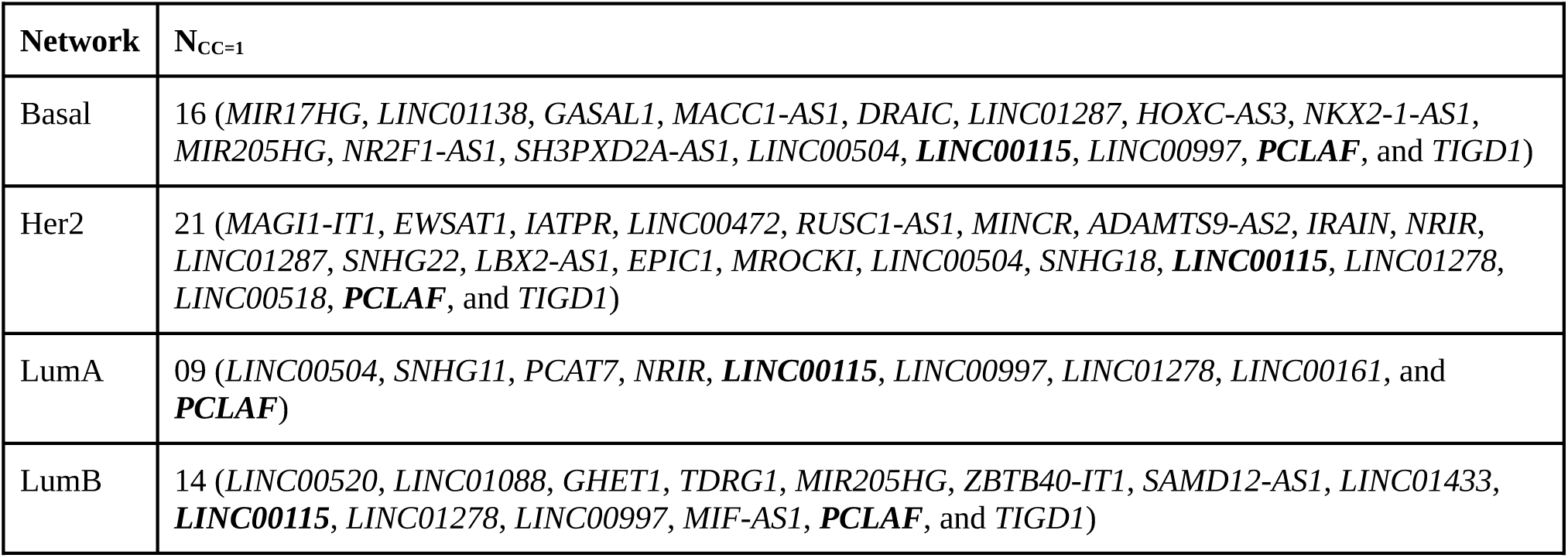
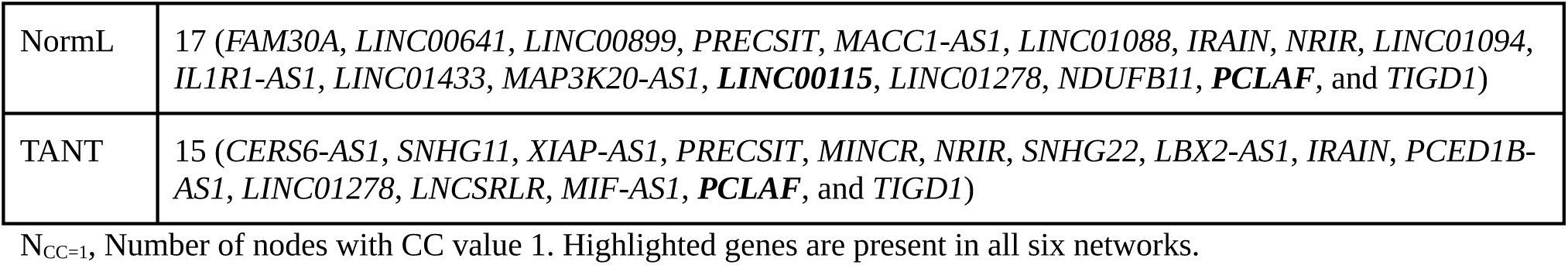
Distribution of clique-forming genes/lncRNAs across all six tissue types.

### Characteristic features of methylated and non-methylated genes

Upon exploring seven different topologies of methylated and non-methylated genes across all six different tissue types, it was interesting to observe that methylated genes exhibit higher average values of degree (Fig. 2), betweenness (Fig. 3), eigenvector centrality (Fig. 6), and pagerank (Fig. 7) compared to non-methylated genes. However, non-methylated genes exhibit higher average values of closeness (Fig. 4) and clustering coefficient (Fig. 5) compared to methylated genes across all six tissue types.

**Fig 2.**
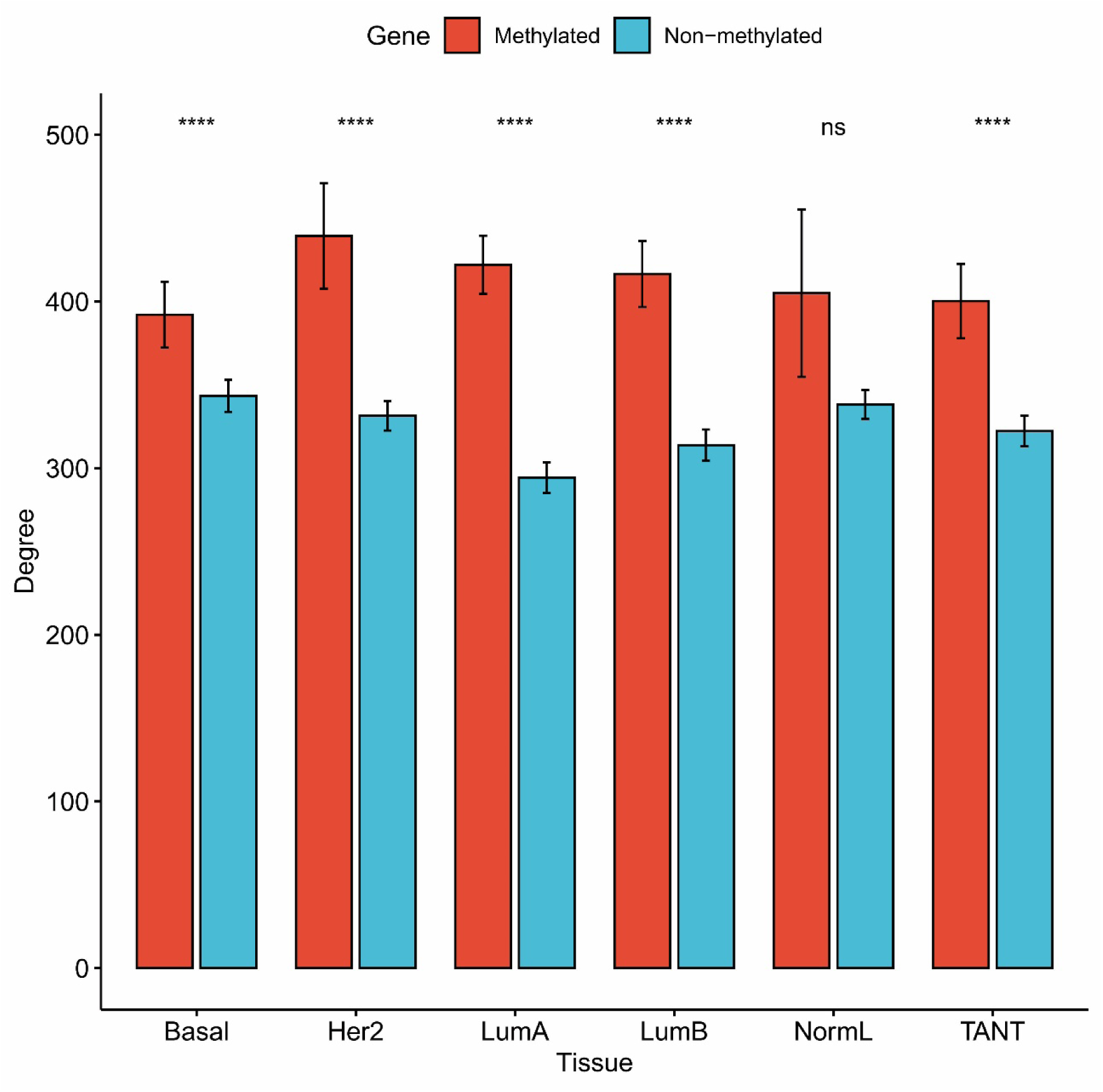
Distribution of degree values across all six tissue types

**Fig 3.**
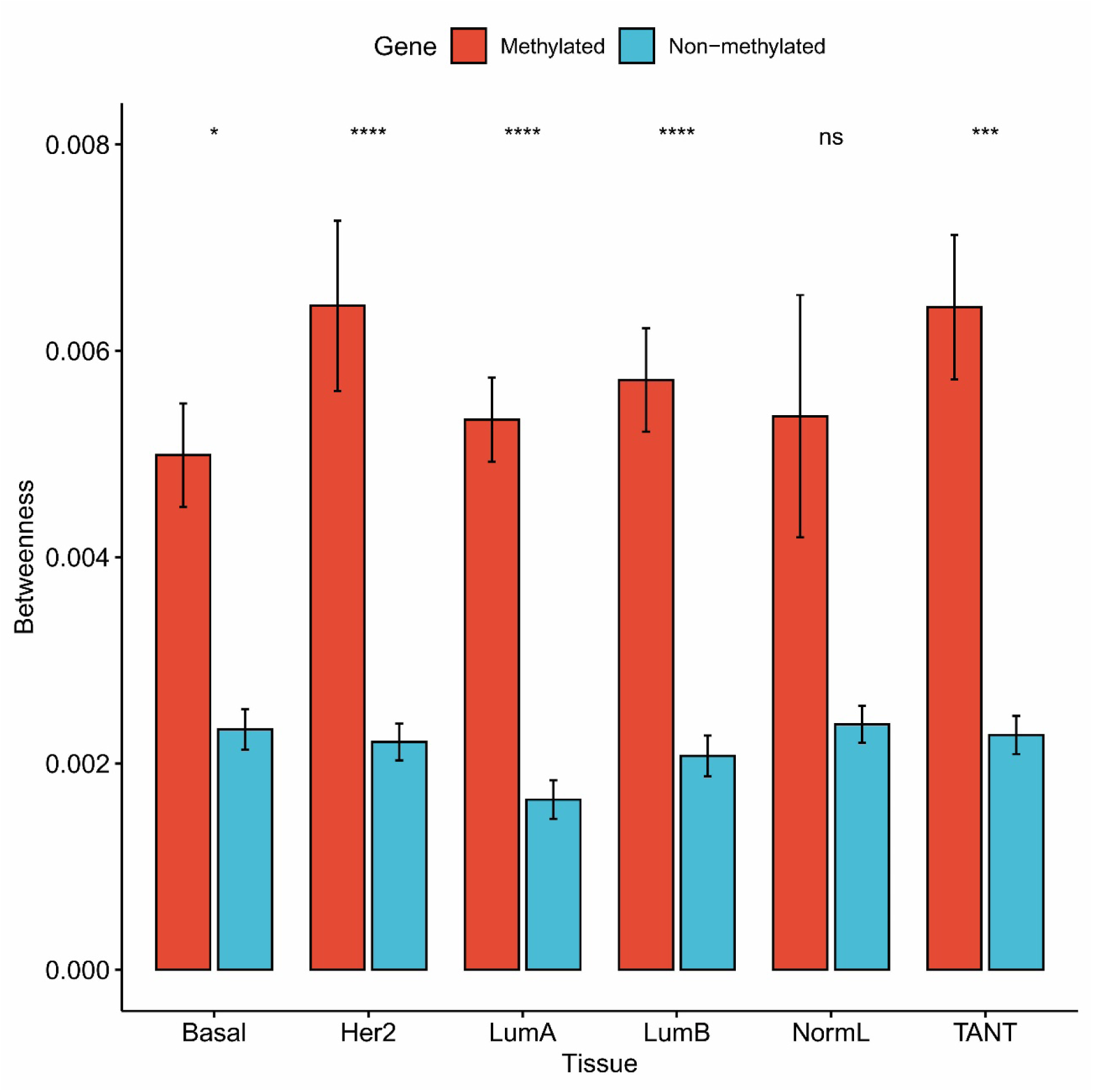
Distribution of betweenness values across all six tissue types

**Fig 4.**
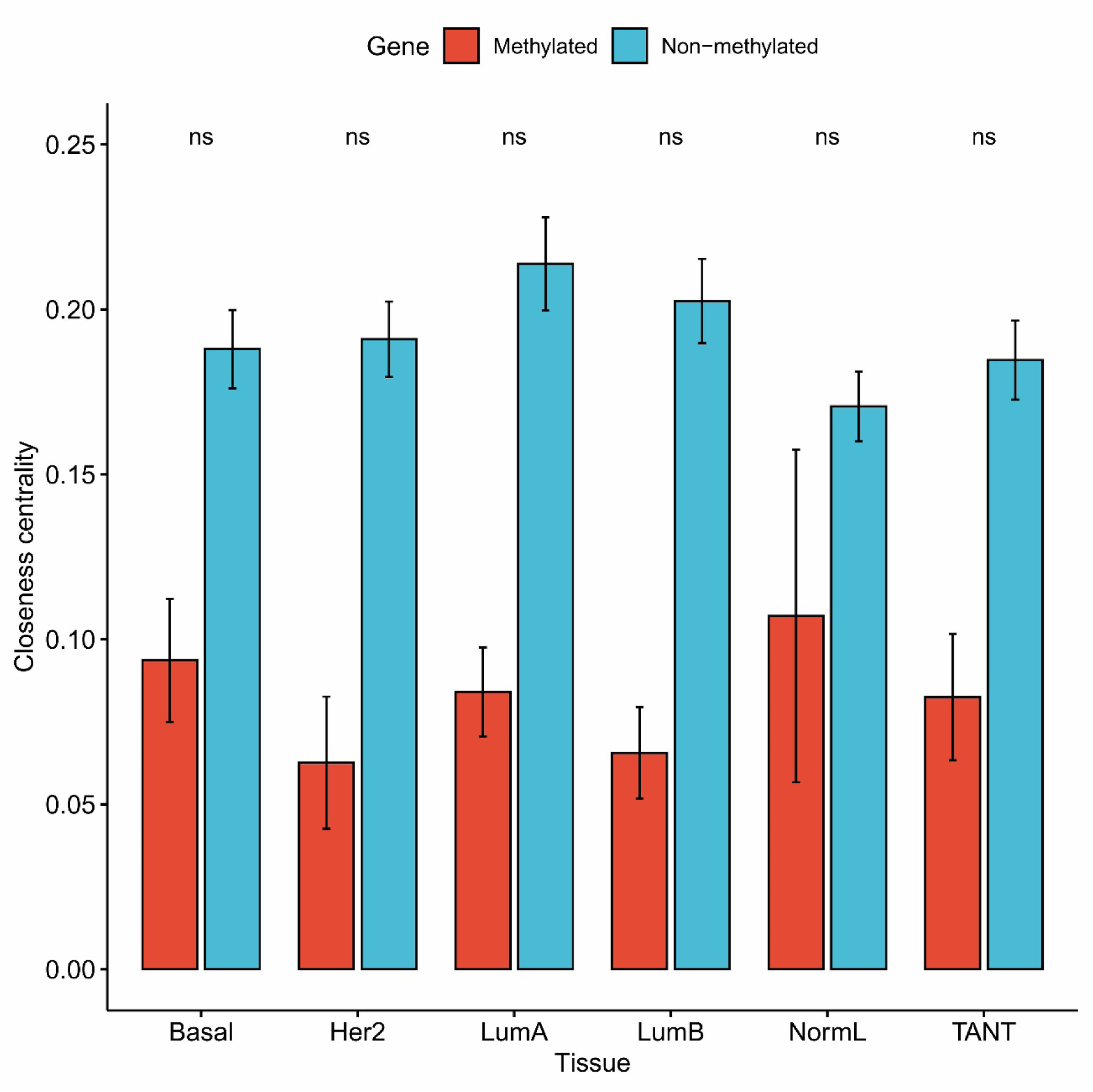
Distribution of closeness values across all six tissue types

**Fig 5.**
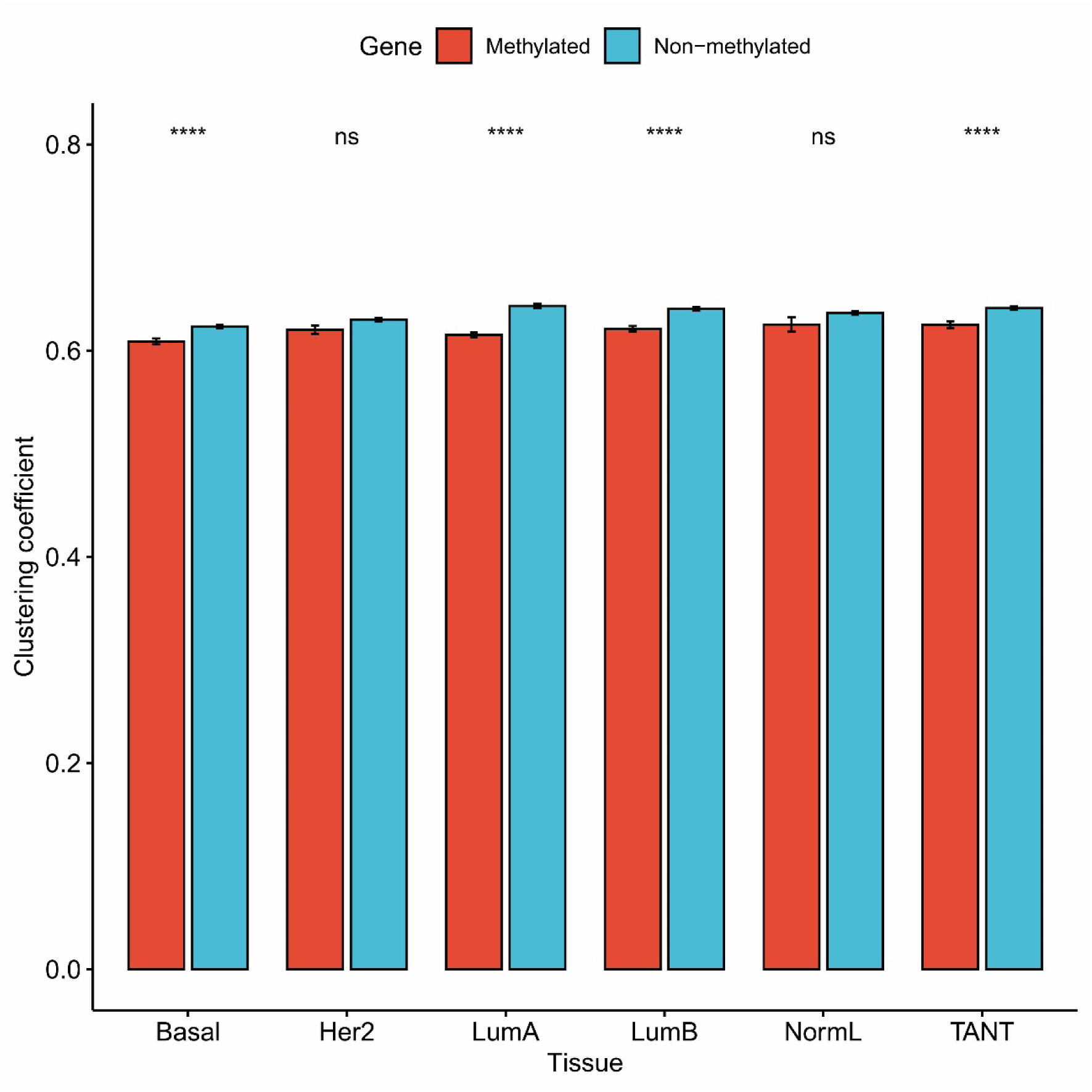
Distribution of clustering coefficient values across all six tissue types

**Fig 6.**
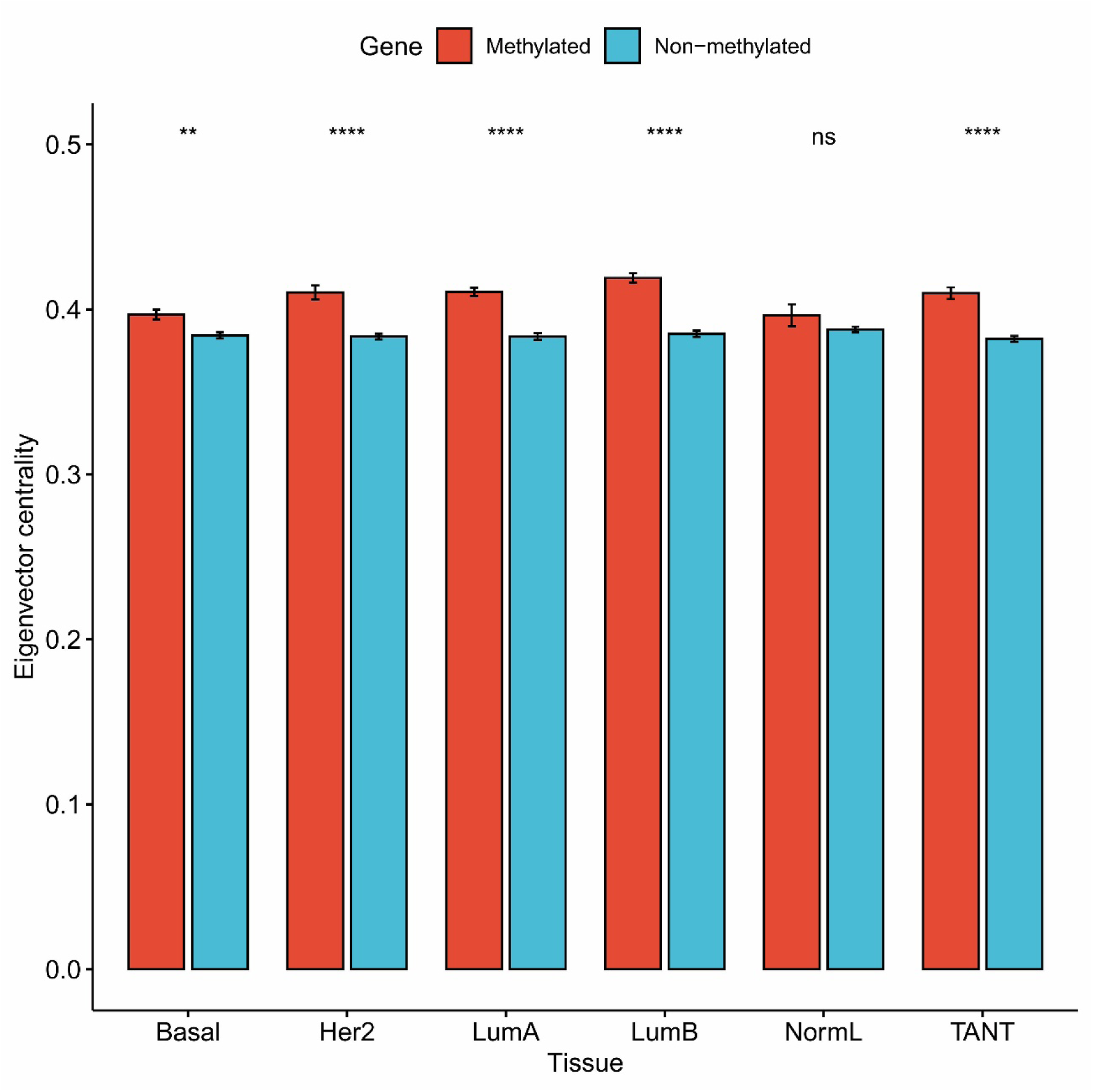
Distribution of eigenvector values across all six tissue types

**Fig 7.**
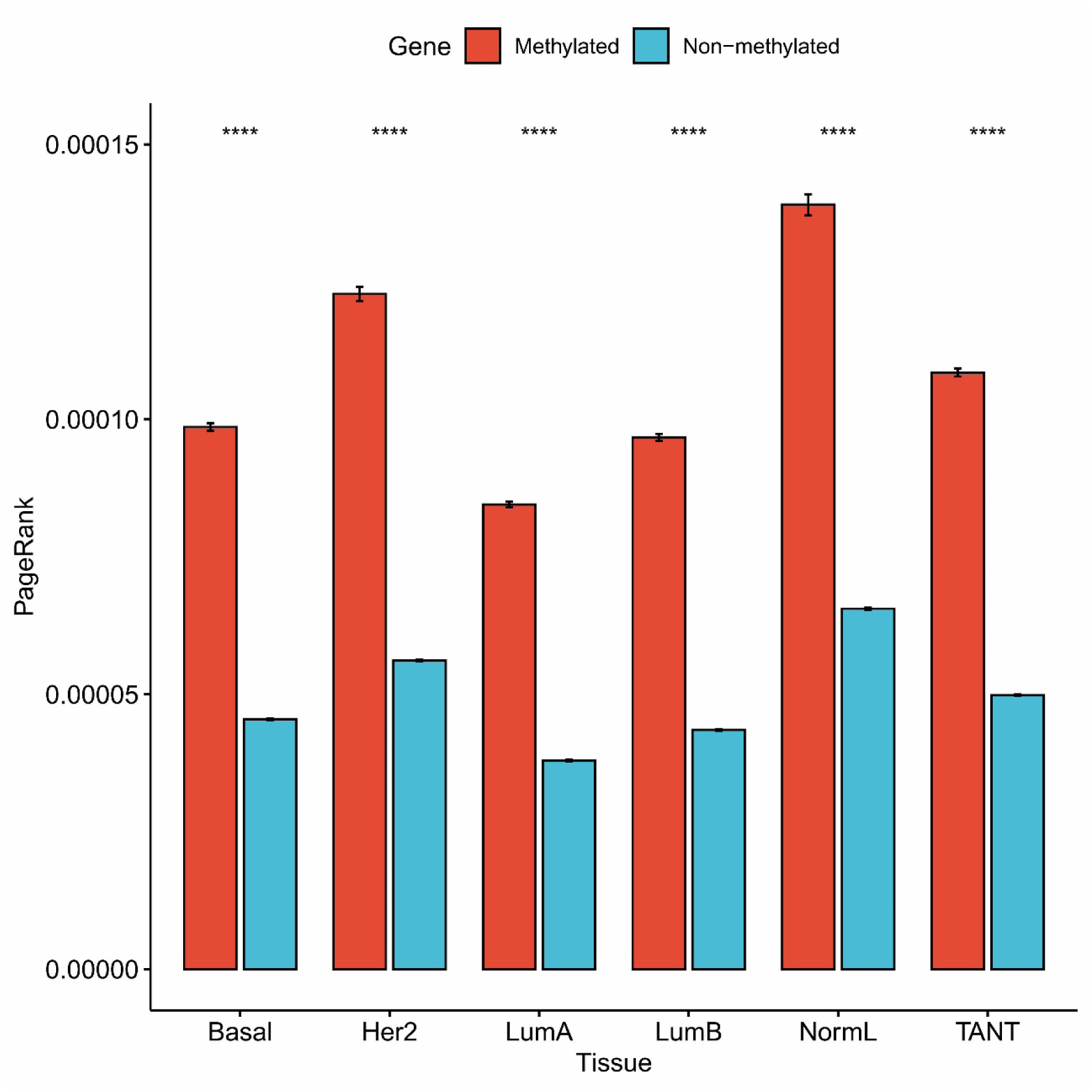
Distribution of pagerank values across all six tissue types

### Pathways associated with hubs and bottlenecks

Pathway enrichment analysis revealed that most hubs and bottlenecks were significantly associated with pathways of human diseases and organismal systems. There were 249 pathways of Human Diseases and 115 pathways of Organismal Systems that were commonly enriched with both hubs and bottlenecks (Tables S4 and S5). Further, among all human disease pathways, we found that the pathways of “Transcriptional misregulation in cancer,” “Human T-cell leukemia virus 1 infection”, “Breast cancer,” “Prostate cancer,” and “Chemical carcinogenesis - receptor activation” were significantly over-represented across all six tissue types.

### Pathway of Transcriptional misregulation in breast cancer

Upon pathway enrichment analysis, we found that the pathway of “Transcriptional misregulation in cancer” was significantly over-represented across all six tissue types. Therefore, to understand this pathway in breast cancer at the epigenetic level across all five subtypes and also in TANTs, we tried to construct and expand the pathway network by mapping it onto the modeled integrative networks of all six tissue types one by one. For this, the pathway “hsa05202” (Transcriptional misregulation in cancer) from the KEGG database, along with five other pathways embedded/linked with it, viz., “hsa04115” (p53 signaling pathway), “hsa05211” (Renal cell carcinoma), “hsa05215” (Prostate cancer), “hsa05216” (Thyroid cancer), and “hsa05221” (Acute myeloid leukemia) were extracted and converted into an edge list using the KEGGgraph [Zhang & Wiemann, 2009] and igraph R package [Csardi & Nepusz, 2006]. This edge list was further used as prior knowledge of known regulations to construct a transcriptional misregulation network (TMRN) for each tissue type by mapping it onto the previously constructed corresponding integrative network of each tissue type. This resulted in six different regulatory pathway networks of Transcriptional misregulation. The common component of TMRN across all six tissue types was investigated and visualized using Cytoscape [Shannon et al. 2003], as available in Fig S1.

## Discussion

Breast cancer is one of the most dreaded and heterogeneous diseases, with the highest number of morbidity and mortality among women worldwide. The limiting factors behind this are mainly the lack of disease awareness, lifestyle, and therapeutic ineffectiveness, including side effects of the current treatment regimen, recurrence of primary tumors, metastasis, and drug resistance. Thus, these factors need to be addressed to effectively manage the disease, which requires an in-depth understanding of the molecular mechanisms of the disease development from a new perspective. Epigenomic analysis coupled with GRNs encompassing both genetic and epigenetic regulations has the immense potential to shed unprecedented insight into the regulatory architecture of cancers and, hence, the molecular mechanisms underlying the development and progression of the disease. In this regard, the current study explores TANTs along with BTTs through a multi-omics perspective involving the construction and analysis of DNAm and ncRNAs-mediated regulatory systems of these tissues. This can lead to the identification of novel diagnostic and therapeutic candidates for the early management of the disease.

When we investigated the methylation pattern of genes and lncRNAs across all six tissue types, it revealed that both genes and lncRNAs were mostly methylated at both promoter and body regions. In contrast, a lesser number of genes and lncRNAs were methylated at their promoter regions only, whereas very few of them were found methylated at their body regions only. Further, some genes were methylated across all tissue types, whereas few were only in the case of some tissue types. For example, *BCL2* and *PART1* were methylated at both body and promoter regions across all six tissue types, whereas *AR* and *HCP5* were methylated at both body and promoter regions across all tissue types except NormL. Moreover, *BRCA1* and *MEG3* were methylated at both body and promoter regions across all tissue types except NormL and TANT. Similarly, some genes were found to be methylated only in TANT but not in other tissue types, such as *TP53* (methylated at promoter region only), *TP63* (methylated at both body and promoter regions), *MAPK12* (methylated at promoter region only), *HCG9* (methylated at body region only), and *LOH12CR2* (methylated at both body and promoter regions).

Among all hubs, 388 protein-coding genes and 100 miRNAs were commonly present as hubs in all six tissue networks, whereas 400 protein-coding genes and 102 miRNAs were commonly present as hubs in all five subtype networks. However, only 15 lncRNAs (viz. *MEG3*, *HCP5*, *UCA1*, *CDKN2B-AS1*, *HOTAIR*, *PVT1*, *GAS5*, *H19*, *XIST*, *DANCR*, *MALAT1*, *SNHG1*, *AFAP1-AS1*, *NEAT1*, and *CLLU1*) were commonly present hubs in all six tissue networks; whereas, 16 lncRNAs (viz. *MEG3*, *HCP5*, *UCA1*, *CDKN2B-AS1*, *HOTAIR*, *PVT1*, *GAS5*, *CRNDE*, *H19*, *XIST*, *DANCR*, *MALAT1*, *SNHG1*, *AFAP1-AS1*, *NEAT1*, and *CLLU1*) were commonly present hubs in all five subtype networks. Similarly, among all bottlenecks, 145 protein-coding genes and three miRNAs were commonly present bottlenecks in all six tissue networks, whereas 177 protein-coding genes and four miRNAs were commonly present bottlenecks in all five subtype networks. Further, only one lncRNA, i.e., *PART1*, was commonly present bottleneck in all six tissue networks. There were 142 protein-coding genes (including some well-known cancer genes, e.g., *AR*, *FOXA1*, *MYC*, *NFKB1*, and *TP63*) and three miRNAs (e.g., hsa-miR-34a, hsa-miR-149, and hsa-miR-141) commonly present as both hubs and bottlenecks (HBs) across all six tissue networks. However, none of the lncRNAs were commonly present as HBs across all six tissue networks. Furthermore, seven protein-coding hub genes (viz., *SPI1*, *STAT5A*, *RORC*, *SPDEF*, *ELF5*, *HOXB2*, and *RUNX3*) out of 388 common protein-coding hub genes and three protein-coding bottleneck genes (viz., *SPI1*, *SPDEF*, and *RUNX3*) out of 145 common protein-coding bottleneck genes, including three protein-coding genes (viz., *SPI1*, *SPDEF*, and *RUNX3*) out of 142 common protein-coding genes as HBs, were methylated across all six tissue types. Six genes, viz., *SPI1*, *RORC*, *SPDEF*, *ELF5*, *HOXB2*, and *RUNX3,* were methylated at both the promoter and body region across all six tissues; however, one gene, i.e., *STAT5A* was methylated only at the promoter region across all six tissues. Furthermore, no methylated hub lncRNAs were commonly present across all six tissues. But, in the case of commonly present bottleneck lncRNAs across all six tissue networks, one bottleneck lncRNA (*PART1*) was methylated. The lncRNA *PART1* was methylated at both the promoter and body region across all six tissue types.

Upon exploration of genes, lncRNAs, and miRNAs with CC, it was found that several genes and lncRNAs across all six tissue networks had CC values one and hence were involved in clique formation. But, none of the miRNAs had a CC value of one. Among all genes and lncRNAs having CC values one, gene *PCLAF* and lncRNA *LINC00115* were present across all five subtypes, whereas only gene *PCLAF* was present across all six tissue networks. The *PCLAF* (PCNA Clamp Associated Factor) gene, also known as *PAF15* or *KIAA0101,* has been previously reported to be overexpressed in breast tumors and was associated with significantly low survival. Additionally, it was reported to interact with BRCA1 and regulate centrosome numbers [Kais et al., 2011]. In another study, it was found that the inhibition of *PCLAF* suppressed the cell cycle progression and cell proliferation by promoting the p53 - Sp1 interaction in breast cancer [Lv et al. 2018]. In the current study, *BRCA1*, *TP53*, and *SP1* were among methylated hub genes across all six tissue networks. The *LINC00115* (Long Intergenic Non-Protein Coding RNA 115) promotes the metastasis of breast cancer by modulating miR-7 and its target *KLF4* [Yuan et al., 2020]. In the present study, this *KLF4* was found as a hub gene across all six tissue networks.

Exploration of different topologies (viz., Degree, Betweenness, Closeness, Clustering coefficient, Eigenvector centrality, and PageRank) of methylated and non-methylated genes and subsequent observations helped us infer their network-based distinct organizational patterns. Genes having a higher number of regulatory relations with other neighboring genes, TFs, or ncRNAs make them preferable for the event of methylation over genes with comparatively lesser numbers of regulatory relations. Further, genes connecting two modules or sub-networks, connecting with genes having higher numbers of neighbors and hence having higher importance than others, were more likely to get methylated than genes not connecting any modules or genes with a higher number of neighbors. Whereas genes closely linked with other genes and the rest of the network and participating in module formation preferably do not undergo methylation. Our study is the first to explore the topological features of methylated genes compared to non-methylated genes in integrative or multilayer networks of six different breast tumor tissues, including TANTs. However, in a brain tissue-based protein-protein interaction network study by Li et al., the topological properties, viz., degree, betweenness, and closeness centrality of genes with low (LMGs) and high methylation (HMGs) have been investigated, and it was observed that the LMGs had a higher average degree, betweenness, and closeness compared to HMGs [Li et al., 2013]. In contrast, the present study includes LMGs and HMGs, both combined as one group of methylated genes. Further, the average closeness centrality values of non-methylated genes were higher than the methylated genes but were not significant (P value > 0.05). This can be attributed mainly to the presence of only a small number of genes with closeness centrality, while a large number of genes had closeness values “NaN” because other genes in the network were not reachable from these genes.

The pathway over-representation or enrichment analysis of hubs and bottlenecks highlighted their involvement mostly in the pathways of disease and organismal systems. Among these, the pathway “Transcriptional misregulation in cancer” was significantly over-represented across all six tissue types, with the highest numbers of both hub and bottleneck genes compared to other pathways. It showed that our multilayer networks of all tissue types, including TANTs, were composed of the sub-network that corresponds to the pathway of “Transcriptional misregulation in cancer” but at the epigenetic level. These sub-networks hold promise to expand our knowledge on the early epigenetic events leading to the development and progression of breast cancer.

In conclusion, we systematically explored multilayered networks of six different tumor-associated tissues of the female breast comprising subtypes and TANTs. These multilayer or integrative networks included four different types of nodes, viz., genes, lncRNAs, miRNAs, and methylation of genes/lncRNAs. The subsequent exploration and analysis of these networks led to the identification of driver candidates such as genes, lncRNAs, and miRNAs, indicating their pivotal roles in the development and progression of the disease. The conserved and varying methylation pattern of genes and lncRNAs across all six tissue types also highlighted their involvement in the disease. Across all six tissue types, genes and lncRNAs were mostly methylated at both the promoter and body regions, followed by the promoter region only and then the body region only. We also mapped the pathway of transcriptional misregulation in cancer at the epigenetic level across all six tissue types, which further can help us understand the early developmental event of the disease. Thus, the findings of this study can lead to the discovery of potential diagnostic and prognostic biomarkers, which could greatly help improve clinical treatment decisions and augment risk prediction.

## Acknowledgments

The authors thank the Center for Modeling, Simulation & Design (CMSD), University of Hyderabad, for providing computational facilities. Further, VV would like to thank the Indian Council of Medical Research (ICMR), New Delhi (ISRM/12(72)/2020, ID: 2020-2951), Institution of Eminence – University of Hyderabad (No. UoH/IoE/RC3-21-052), and Department of Biotechnology (DBT), Government of India (No. BUILDER-DBT-BT/INF/22/SP41176/2020), for their financial support. SK also acknowledges the Institution of Eminence – UoH for the Performance-Based Publication Incentives, University of Hyderabad for the Non-NET Fellowship, and ICMR for the Senior Research Fellowship (Grant No.: 3/2/2/113/2019/NCD-III, ID: 2019-6723).

## Author Contributions

SK and VV conceived and designed the study. SK performed the experiments and analyzed the data. SK drafted, wrote, and edited the manuscript. VV reviewed and edited the manuscript. VV supervised the study. All authors read and approved the final manuscript.

