## Supplementary Fig. S1 for "Decoding DNA methylation and non-coding RNAs mediated regulatory landscape of breast tumor and adjacent normal tissues"

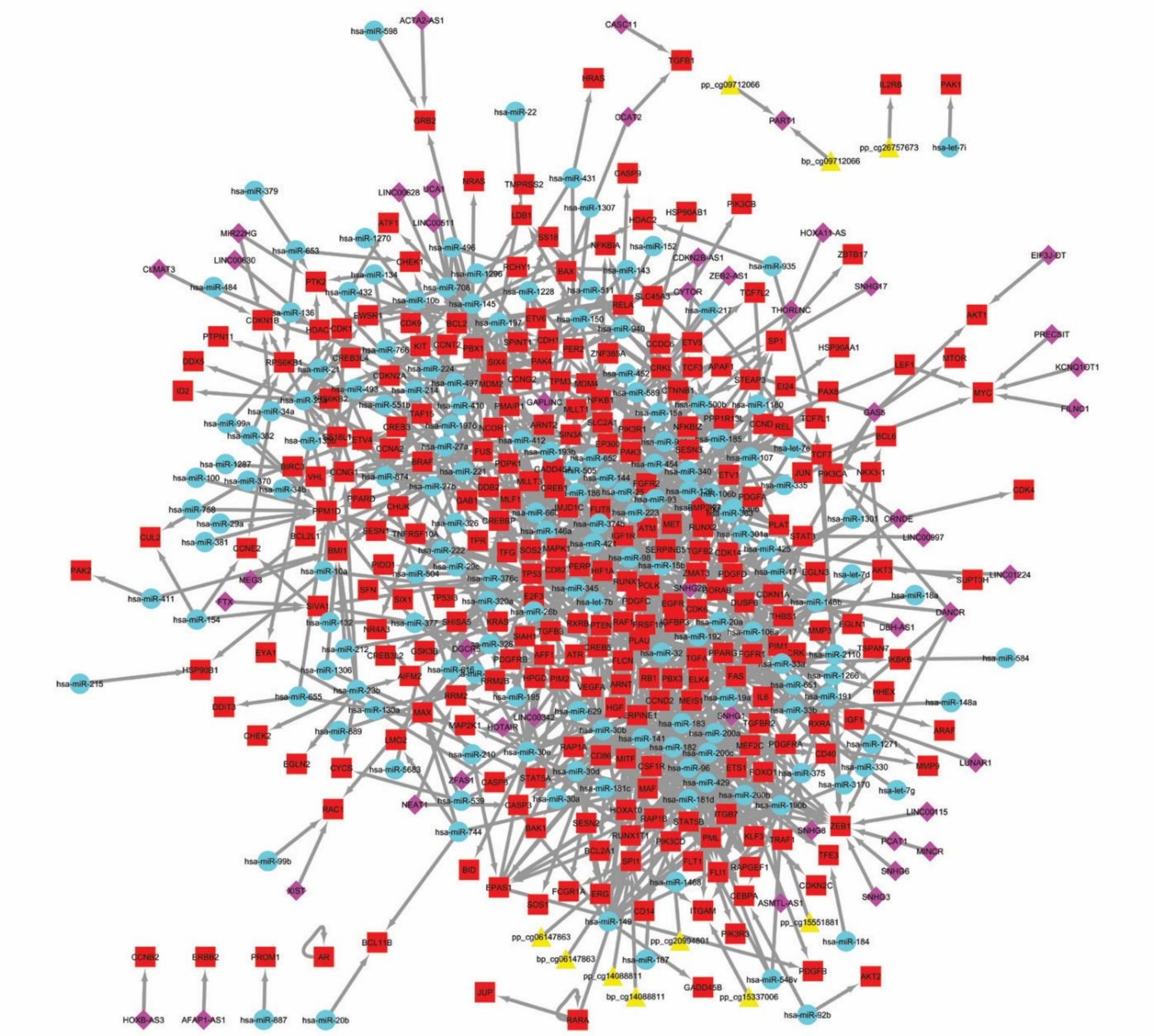

**Fig S1**. Sub-network representing transcriptional misregulation commonly present across all six tissue networks
