## Supplementary Tables S1, S2, and S3 for "Decoding DNA methylation and non-coding RNAs mediated regulatory landscape of breast tumor and adjacent normal tissues"

### Supplementary tables S1 – S3

Table S1. Distribution of hub genes and miRNAs across all six tissue types.

| Network | **Hubs genes** | **Hub miRNAs** | **Methylated** | **Common Hubs (Genes)** | **Common Hubs (miRNAs)** | **Common Hubs (Methylated)** |
|---|---|---|---|---|---|---|
| Basal | 471 | 146 | 127 | 388 | 100 | 7 (*RUNX3*, *RORC*, *SPDEF*, *ELF5*, *SPI1*, *HOXB2*, *STAT5A*) |
| Her2 | 449 | 131 | 81 | | | |
| LumA | 443 | 141 | 225 | | | |
| LumB | 440 | 141 | 161 | | | |
| NormL | 441 | 117 | 23 | | | |
| TANT | 445 | 128 | 113 | | | |

Table S2. Distribution of bottleneck genes and miRNAs across all six tissue types.

| Network | **Bottleneck genes** | **Bottleneck miRNAs** | **Methylated** | **Common Bottlenecks (Genes)** | **Common Bottlenecks (miRNAs)** | **Common Bottlenecks (Methylated)** |
|---|---|---|---|---|---|---|
| Basal | 335 | 19 | 5 | 145 | 3 (hsa-miR-34a, hsa-miR-149, hsa-miR-141) | 3 (*SPI1*, *SPDEF*, *RUNX3)* |
| Her2 | 341 | 16 | 6 | | | |
| LumA | 323 | 21 | 5 | | | |
| LumB | 317 | 13 | 5 | | | |
| NormL | 324 | 23 | 5 | | | |
| TANT | 302 | 20 | 6 | | | |

Table S3. Distribution of hub and bottleneck lncRNAs across all six tissue types.

| Network | **Hubs** | **Common Hubs** | **Bottlenecks** | **Common Bottlenecks** |
|---|---|---|---|---|
| Basal | 23 (MEG3, HCP5, SNHG3, KCNQ1OT1, UCA1, CDKN2B-AS1, HOTAIR, PVT1, GAS5, CRNDE, H19, XIST, DANCR, | 15 (MEG3, HCP5, UCA1, CDKN2B-AS1, HOTAIR, PVT1, GAS5, H19, XIST, DANCR, MALAT1, SNHG1, AFAP1-AS1, NEAT1, and CLLU1) | 7 (MEG3, PART1, HCP5, SNHG3, KCNQ1OT1, SNHG4, and NBR2) | 1 (PART1) |

| | | | |
|---|---|---|---|
| | MALAT1, HOXA11-AS, SNHG1, AFAP1-AS1, SNHG6, NEAT1, CYTOR, DIRC1, CT62, and CLLU1) | | |
| Her2 | 21 (PVT1, MEG3, HCP5, H19, CDKN2B-AS1, GAS5, CRNDE, XIST, UCA1, DANCR, MALAT1, HOXA11-AS, HOTAIR, ZFAS1, CYTOR, AFAP1-AS1, NEAT1, SNHG1, OIP5-AS1, CT62, and CLLU1) | | 7 (PVT1, MEG3, PART1, HCP5, KCNQ1OT1, SNHG4, and NBR2) |
| LumA | 21 (H19, PVT1, MEG3, CASC2, SNHG1, HCP5, HOXA11-AS, HOTAIR, CRNDE, XIST, UCA1, DANCR, CYTOR, MALAT1, CDKN2B-AS1, GAS5, AFAP1-AS1, NEAT1, DIRC1, CT62, and CLLU1) | | 12 (H19, PVT1, MEG3, CASC2, ATXN8OS, SNHG1, SNHG3, KCNQ1OT1, PART1, HCP5, NBR2, and MIR17HG) |
| LumB | 21 (MEG3, KCNQ1OT1, HCP5, H19, HOTAIR, PVT1, CRNDE, XIST, UCA1, DANCR, CDKN2B-AS1, SNHG12, SNHG1, AFAP1-AS1, MALAT1, CASC2, GAS5, NEAT1, DIRC1, CT62, and CLLU1) | | 7 (MEG3, ATXN8OS, PART1, KCNQ1OT1, SNHG4, HCP5, and NBR2) |
| NormL | 18 (H19, CDKN2B-AS1, HOXA11-AS, HOTAIR, GAS5, CRNDE, PVT1, MALAT1, MEG3, UCA1, CYTOR, AFAP1-AS1, DANCR, XIST, NEAT1, SNHG1, HCP5, and CLLU1) | | 3 (H19, PART1, and BCYRN1) |
| TANT | 21 (PVT1, H19, SNHG1, HCP5, NEAT1, CDKN2B-AS1, HOXA11-AS, HOTAIR, XIST, | | 7 (PVT1, H19, SNHG1, HCP5, PART1, NBR2, and EMX2OS) |

| | |
|---|---|
| | UCA1, DANCR, MALAT1, MEG3, GAS5, CYTOR, AFAP1-AS1, MIAT, SNHG16, KCNQ1OT1, CLLU1, and PLAC4) |
